# Actomyosin-mediated nanostructural remodeling of the presynaptic vesicle pool by cannabinoids induces long-term depression

**DOI:** 10.1101/444950

**Authors:** Maureen H. McFadden, Hao Xu, Yihui Cui, Rebecca A. Piskorowski, Christophe Leterrier, Diana Zala, Laurent Venance, Vivien Chevaleyre, Zsolt Lenkei

**Author notes:** co-first authors (Maureen H. McFadden, Hao Xu). co-senior authors (Laurent Venance, Vivien Chevaleyre, Zsolt Lenkei). **Corresponding author** Zsolt Lenkei, Center of Neuroscience and Psychiatry, INSERM U894, rue de la Santé 102-108, 75015, Paris, France.

## Abstract

Endo- and exocannabinoids, such as the psychoactive component of marijuana, exert their effects on brain function by inducing several forms of synaptic plasticity through the modulation of presynaptic vesicle release^1-3^. However, the molecular mechanisms underlying the widely expressed endocannabinoid-mediated long-term depression^3^ (eCB-LTD), are poorly understood. Here, we reveal that eCB-LTD depends on the contractile properties of the pre-synaptic actomyosin cytoskeleton. Preventing this contractility, both directly by inhibiting non-muscle myosin II NMII ATPase and indirectly by inhibiting the upstream Rho-associated kinase ROCK, abolished long-term, but not short-term forms of cannabinoid-induced functional plasticity in both inhibitory hippocampal and excitatory cortico-striatal synapses. Furthermore, using 3D superresolution microscopy, we find an actomyosin contractility-dependent redistribution of synaptic vesicle pools within the presynaptic compartment following cannabinoid receptor activation, leading to vesicle clustering and depletion from the pre-synaptic active zone. These results suggest that cannabinoid-induced functional plasticity is mediated by a nanoscale structural reorganization of the presynaptic compartment produced by actomyosin contraction. By introducing the contractile NMII as an important actin binding/structuring protein in the dynamic regulation of synaptic function, our results open new perspectives in the understanding of mechanisms of synaptic and cognitive function, marijuana intoxication and psychiatric pathogenesis.

## Introduction

Brain connectivity patterns arise through protracted developmental events that lead to assembly, activity-based selection and stabilization of synapses in the mature brain, through precise regulation of cytoskeletal dynamics. Importantly, synapses retain important functional plasticity, a critical component for experience-dependent adjustments of brain function. Both axons and presynaptic terminals undergo long-term experience and activity-dependent structural plasticity^4^, similar to postsynaptic dendritic spines, but molecular mechanisms are not well known. Several established and widespread forms of functional presynaptic plasticity throughout the mammalian brain are retrograde and endocannabinoid-mediated^1-3^. Activity-dependent release of endocannabinoids (eCBs) by the postsynaptic cell, followed by the activation of the presynaptic type-1 cannabinoid receptor (CB_1_R), mediates either a short-term depression (STD) of transmitter release, such as depolarization-induced suppression of inhibition (DSI) or excitation (DSE), or a long-term plasticity, such as eCB-mediated long-term depression (eCB-LTD)^3^. Although the mechanisms underlying DSI and DSE have been well established, the presynaptic mechanisms mediating eCB-LTD, much like other forms of long-term presynaptic plasticity, remain poorly understood.

Recently, we described a novel molecular mechanism through which the type-1 cannabinoid receptor (CB_1_R), an abundant and well-known mediator of synaptic plasticity in the adult brain, exerts neurodevelopmental effects in the embryonic brain^5^. In this mechanism, endo- and exo-cannabinoids, such as Δ^9^THC, the psychoactive compound of marijuana, act through CB1Rs to induce non-muscle myosin II (NMII) contractility, downstream of the small the GTPase RhoA and Rho - associated kinase (ROCK). This leads to axonal growth cone retraction and to rapid and lasting changes in the morphology of developing neurons^5^. Because mediators of neuronal development and neurite growth have been proposed to retain their structural roles in the mature brain, albeit on a smaller spatial scale^6^, we investigated here the possible involvement of this molecular mechanism in CB_1_R functions regulating synaptic plasticity in the mature brain by combining single-cell patch-clamp recordings with a novel quantitative analysis of presynaptic nanoarchitecture.

## Results and Discussion

First, we inhibited actomyosin contractility in acute brain slices from two different brain regions (**Fig. 1a, 2a**), during well-established forms of eCB-mediated STD and LTD, by preincubating slices for 20 minutes with blebbistatin (10μM), a selective NMII ATPase inhibitor^7^. In hippocampal slices of 6-8 weeks old mice, DSI, a STD of GABA release triggered by depolarization of pyramidal cells, was not affected by this treatment (**Fig.1b**); however, application of blebbistatin, but not of its inactive enantiomer, strikingly abolished the eCB-mediated LTD of inhibitory post-synaptic currents (IPSCs) (**Fig. 1c**). ROCK inhibition with the selective inhibitor Y-27632 (10μM) also fully blocked eCB-LTD induction (**Fig. 1h**). Blebbistatin application also abolished the change in release probability following eCB-LTD induction (as indirectly measured by the paired-pulse ratio PPR) (**Fig. 1d**), but did not affect the initial PPR (**Fig. 1d)**. This indicates that blebbistatin prevented the eCB-mediated decrease in presynaptic vesicle release without altering basal transmission. In the hippocampus, induction of eCB-LTD requires not only CB_1_R activation, but also spontaneous firing of GABAergic interneurons. We therefore tested whether blebbistatin might indirectly alter eCB-LTD by reducing interneuron firing. We found that blebbistatin did not change the amplitude or the frequency of spontaneous IPSCs (**Fig. 1e.**). Furthermore, by applying a protocol capable of rescuing eCB-LTD when interneuron firing is blocked, using trains of stimulation following tetanus^8^, we found that eCB-LTD was still not induced in the presence of blebbistatin (**Fig. 1f.**). These data strongly indicate that blebbistatin is acting at presynaptic inhibitory terminals and is not altering interneuron activity. Finally, because eCB-LTD induction requires glutamate release, mGluR activation and eCB release, we bypassed all these steps and directly looked at the effect of CB_1_R activation with the high affinity agonist WIN55,212-2^9^ on action potential-independent IPSCs, i.e. miniature IPSCs (mIPSCs). We found that WIN55,212-2 induced a significant decrease in the frequency but not the amplitude of mIPSCs, confirming the presynaptic origin of CB_1_R activation (**Fig. 1g**). Strikingly, in the presence of blebbistatin, WIN55,212-2 had no effect on mIPSC frequency (**Fig. 1g**).

**Figure 1:**
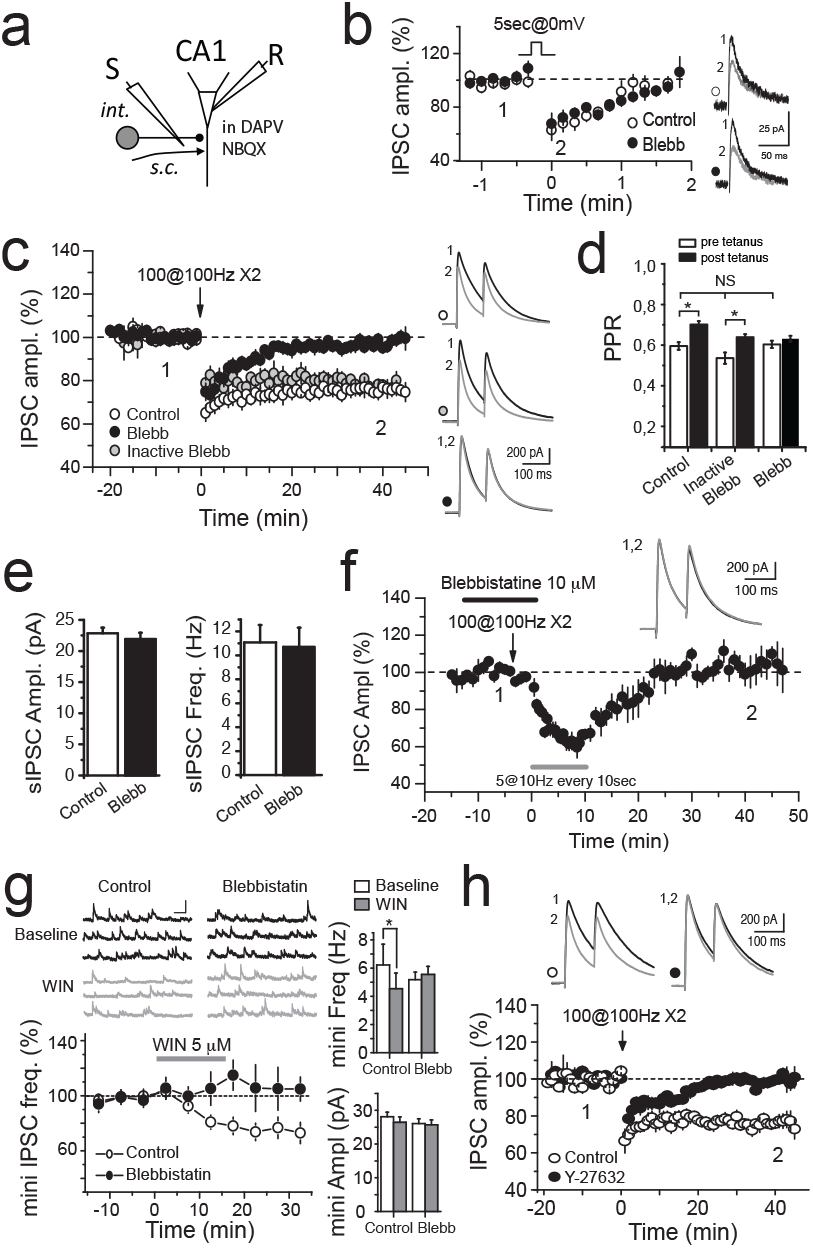
Presynaptic actomyosin contractility mediates eCB-LTD at inhibitory synapses in the CA1 area of the hippocampus. (**a**) Schematic of recording paradigm. CA1: whole-cell recorded pyramidal neuron; int: inhibitory interneuron fibers, activated by a stimulating electrode in the stratum radiatum. Fast excitatory transmission from Schaffer collateral inputs (s.c.) was blocked by the AMPA/NMDA/KA receptor antagonists D-APV and NBQX. (**b**) The transient depression of inhibitory transmission following a 5 second depolarization at 0 mV (white circles) was unaffected by blebbistatin (Blebb, 10μM, black circles). In control: 65.2±6.5% of baseline, n=7. In blebbistatin: 69±5% of baseline, n=6, p=0.63. (**c**) The long-term depression following high frequency stimulation (white circles) was completely abolished in the presence of blebbistatin (black circles) but not by the inactive blebbistatin enantiomer (10μM, grey circles). Average sample traces are shown on top for time points (1) and (2). Control: 75±3% of baseline, n=9; blebbistatin: 97±2%, n=8, p=0.00002, inactive blebbistatin 79±2%, n=5, p=0.27. **(d)** Tetanic stimulation resulted in a significant increase (p = 0.003) in the paired pulse ratio (PPR), in accordance with the decrease in GABA release during LTD. This increase in PPR was not affected in presence of inactive blebbistatin but was abolished in presence of blebbistatin. Note that Blebbistatin did not induced any change in PPR before the tetanus (compared to the initial PPR in control), indicating that basal release probability was not altered by Blebbistatin. **(e)** Blebbistatin did not change the amplitude or the frequency of spontaneous IPSCs. Amplitude: baseline: 22.8 ±0.9pA, after blebbistatin: 21.9±1.0pA, n=7, p=0.18; frequency: baseline: 11.1±1.4Hz, after blebbistatin: 10.7±1.6Hz, p=0.59. **(f)** Trains of stimulation following tetanus do not result in eCB-LTD in the presence of blebbistatin. 104±6% of baseline, n=5. (**g**) The decrease in miniature IPSC frequency mediated by WIN55,212-2 (WIN, 5μM, white circles) was abolished by blebbistatin (black circles). Sample traces are shown on top. Right: Average mIPSC frequencies and amplitudes. Control: 1.22±0.03 of baseline, p = 0.0029 Frequency: from 6.2±1.4 to 4.5±1.1Hz, p=0.03; Amplitude: from 28.±1.4 to 26.5±1.6pA, p=0.46, n=5; blebbistatin: 1.05±0.01%, p=0.08, Frequency: from 5.2±0.5 to 5.6±0.6Hz, p=0.6; inactive blebbistatin: 1.21±0.04 p=0.02. (**h**) The LTD evoked by high frequency stimulation (white circles) was abolished in the presence of the ROCK inhibitor Y-27632 (10μM, black circles). Average sample traces are shown on top for time points (1) and (2). 99±1% of baseline, n=5, p=0.37 with baseline and p=0.0004 with control LTD, n=6. Student’s t-test. Values: mean± SEM.

While in the hippocampus eCB-induced plasticity is mostly present at inhibitory synapses, it is also widely expressed at excitatory synapses throughout the brain^1-3^. We therefore investigated the molecular mechanism of eCB-STD and LTD at an excitatory glutamatergic synapse: the corticostriatal synapse at medium-sized spiny neurons (MSNs) of the dorsolateral striatum (**Fig. 2a**) of young rats, which expresses eCB-mediated and CB_1_R-dependent STD, namely a DSE, and LTD^10,11^. Here, a sustained depolarization of MSNs induced a DSE (**Fig. 2b**), which, similar to hippocampal DSI, was not significantly affected by blebbistatin (10μM) treatment (**Fig. 2b**). We then tested the striatal eCB-LTD induced after cortical low frequency stimulation (LFS)^10, 12^. This LTD was indeed CB_1_R-mediated as it was prevented by treatment with the CB_1_R specific inhibitor AM251 (3μM) (**Fig. 2c**). As for hippocampal synapses, actomyosin contraction was found to be necessary for corticostriatal eCB-LTD, which was prevented by treatment with blebbistatin, but not with the inactive blebbistatin enantiomer (**Fig. 2d**). This effect was not due to alterations in basal transmission as we found no significant change in PPR for 50ms intervals inter-stimuli in any tested condition (**Fig. 2e**). We further evaluated whether the effect of blebbistatin on eCB-LTD was pre- or postsynaptic by measuring the PPF ratio before and after LFS (PPF_plasticity/baseline_). We found a significant increase in PPF in control conditions whereas no significant variation of PPF was found following treatment with active blebbistatin (**Fig. 3f**), indicating presynaptic action of actomyosin contraction under eCB-LTD. The specific ROCK inhibitor Y-27632 (10μM) also impaired eCB-LTD induction (**Fig. 2g**). Therefore, similarly to hippocampal GABAergic synapses, activation of CB_1_R by eCBs in corticostriatal excitatory synapses induces LTD, but not STD, through ROCK-mediated presynaptic actomyosin contraction.

**Figure 2:**
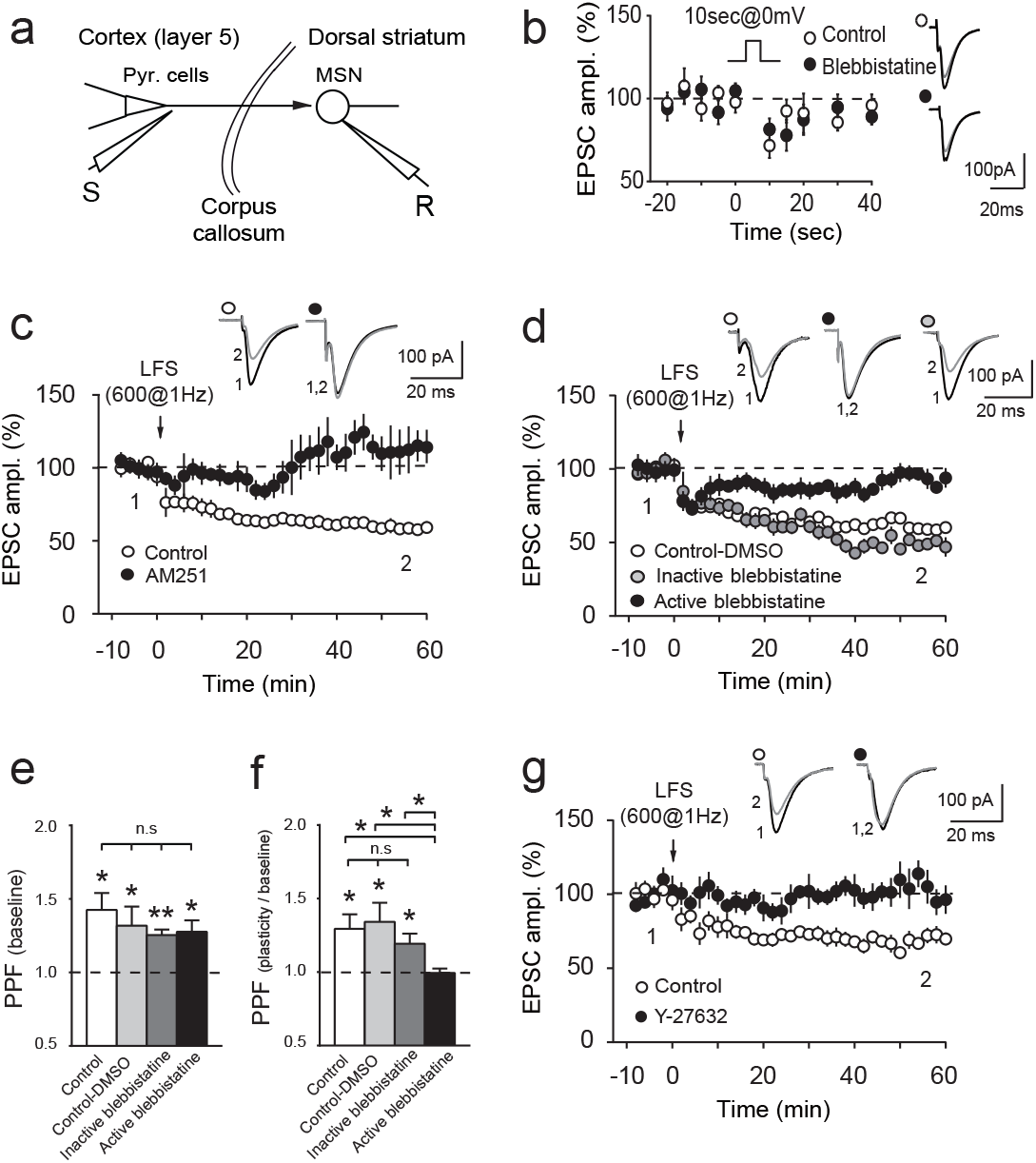
Presynaptic actomyosin contractility mediates eCB-LTD at excitatory corticostriatal synapses. (**a**) Schematic of whole-cell recording of a MSN and stimulation in the somatosensory cortical layer V. MSN: medium-sized spiny neurons (**b**) The transient depression of excitatory transmission following a 10 second depolarization at 0mV (white circles, 81±4% of baseline, n=13, p=0.0003) was unaffected by blebbistatin (black circles, 79±3% of baseline, n=11, p=0.0001 with baseline; p=0.8039 with control DSE, n=13). Average sample traces before and 10sec after the depolarization are shown on the right. (**c**) LTD induced with LFS (control: 58±2% of baseline, n=7, p<0.0001 with baseline) was CB_1_R-mediated because prevented with AM251 (107±10% of baseline, n=5, p=0.5244 with baseline, p<0.0001 with control). (**d**) The eCB-LTD following a LFS (white circles, control-DMSO: 56±7% of baseline, n=9, p=0.0002, p=0.8217 with control without DMSO) was abolished in the presence of blebbistatin (black circles, 95±3% of baseline, n=10, p=0.1201 with baseline, p<0.0001 with control LTD) but was unaffected by the inactive enantiomer of blebbistatin (grey circles, 10μM, 52±4% of baseline, n=11, p<0.0001 with baseline, p=0.0556 with control LTD). Average sample traces are shown on top at the time point before (1) and after the stimulation protocol (2). (**e**) 50ms inter-stimuli intervals induced significant PPR in control (p=0.0203, n=5), control-DMSO (p=0.0120, n=8), inactive blebbistatin (p=0.0023, n=5) and active blebbistatin (p=0.0375, n=4) conditions (Anova: p=0.6291 and F(3, 18)=0.5905). (**f**) PPR_plasticity/baseline_ displayed significant increase in control (PPR_plasticity/baseline_=1.30±0.10, p=0.0397, n=5) control-DMSO: inactive blebbistatin: control, control-DMSO (PPR_plasticity/baseline_=1.34±0.13, p=0.0335, n=8) and inactive blebbistatin (PPR_plasticity/baseline_=1.19±0.07, p=0.0493, n=5; Anova: p=0.6630 and F(2, 15)=0.4225) but not for active blebbistatin (PPR_plasticity/baseline_=1.00±0.03, p=0.8697, n=4). (**g**) The ROCK inhibitor Y-27632 abolishes the eCB-LTD (black circles, 98±11% of baseline, n=7, p=0.7204 with baseline, p=0.0038 with control LTD, n=6). Average sample traces are shown on top for time points (1) and (2). Student’s t-test. Values: mean± SEM.

As blebbistatin directly inhibits the contractility of the actomyosin cytoskeleton, the effects described above suggest that cannabinoid-mediated LTD may be elicited through actomyosin-contractility-induced cytoskeletal remodeling of the presynaptic compartment. In order to evaluate this hypothesis, we chose to directly image the nanoscale presynaptic architecture in dissociated cultures of rat hippocampal neurons. First, we confirmed that actomyosin contractility is involved in cannabinoid-induced synaptic plasticity in this experimental model by measuring the effect of CB_1_R activation on synaptic vesicle release at individual axonal boutons^13^, by using synaptophysin-pHluorin (SpH), which increases in green fluorescence intensity upon vesicle fusion^14^. Neuronal depolarization (KCl; 2min; 50mM) induced an average release of around 30% of total bouton vesicle pool under control conditions (**Fig. 3**), as estimated through maximal bouton fluorescence upon terminal alkaline incubation (NH_4_Cl; 2min, 50mM). Importantly, vesicle release under WIN55,212-2 (1μM; 10min) was significantly lower as compared to control (**Fig. 3**) and this effect was prevented both by pretreating neurons with para-nitroblebbistatin (25μM, **Fig. 3c**), a C15 derivative of blebbistatin^7^ with reduced blue-light sensitivity^15^ and neuronal cytotoxicity (Extended Data Fig. 1), as well as through pretreatment with Y-27632 (10μM, **Fig. 3c**). Therefore, as in acute slices, activation of endogenous CB_1_R decreases vesicle release via ROCK-mediated NMII activation at individual axonal boutons *in vitro*.

**Figure 3:**
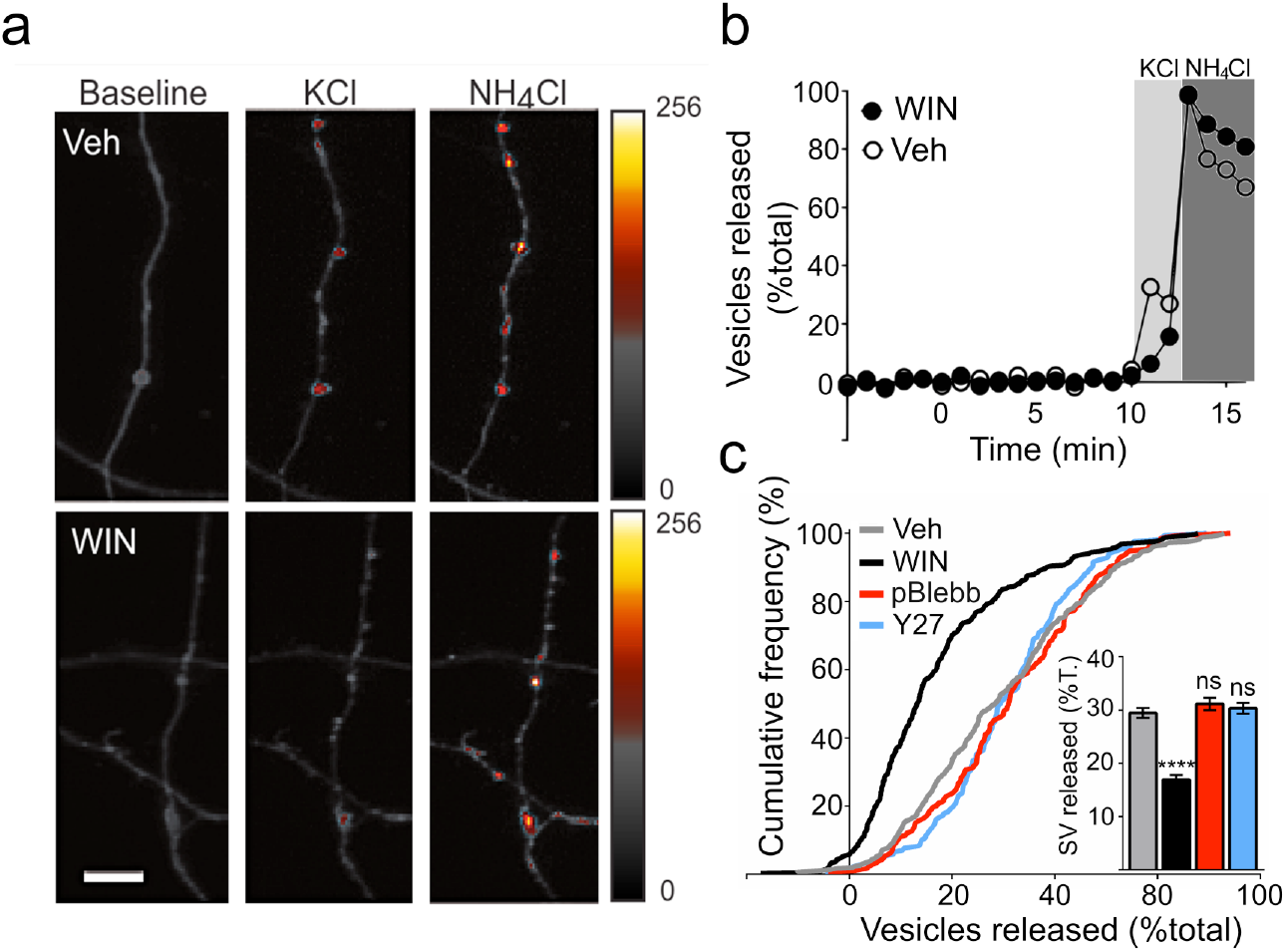
Actomyosin contractility mediates cannabinoid-induced suppression of vesicle release, as shown by pretreatment with the NMII inhibitor para-nitroblebbistatin (pBlebb), or the ROCK inhibitor Y-27632 (Y27). (**a**) Neuron expressing the vesicle release marker SpH. Example SpH fluorescence levels in a control (Veh) and a WIN55,212-2-treated (WIN, 1μM; 10min) axon before stimulation (Baseline), after stimulation (KCL, 50mM, 2min), and after terminal superfusion with NH_4_Cl (50mM, 2min). Fluorescence intensity (arbitrary units) increases during stimulation in control conditions while WIN decreases this effect. (**b**) Experimental paradigm and example traces of normalized axonal bouton SpH fluorescence (**c**) Cumulative probability distributions of the released vesicle pool fractions under control conditions (±0.92%; n=337 over 4 independent experiments), or after treatment with WIN: (16.94%±0.84%; n*=*323 over 4 independent experiments; *P<*0.0001), pBlebb+WIN: (pBlebb: 25μM; 20min; 31.14%±1.15%; n=173 over 3 independent experiments; *P*<0.0001); Y-27632+WIN (Y27: 10μM; 20min; 30.35%±1.03%; n=168 over 3 independent experiments; *P*<0.0001;. ****: p < 0.001); ns: not significant as compared to vehicle, Kruskal-Wallis test. Scale bar: 5μm

We next assessed the nanoarchitecture of the presynaptic compartment and the potential influence of actomyosin dynamics by testing the spatial relationship between synaptic vesicles and the synaptic active zone (AZ) by using the activator/reporter pairing method of multicolor 3D Stochastic Optical Reconstruction Microscopy (STORM)^16^, which allows imaging proteins of interest simultaneously through the same optical path, reducing spatial differences that may arise through optical aberrations. Furthermore, to avoid any artifacts resulting from cross-talk of closely localized fluorophores, we used as a closely apposed but spatially separated spatial AZ reference Homer1, a major protein of the postsynaptic scaffold (PSD, **Fig. 4a**). As pre- and post-synaptic compartments have to be precisely aligned inside individual synaptic nanomodules^17^ in order to ensure sensitive and efficient detection of synaptic events, AZ and PSD synaptic scaffoldings are highly correlated^17,18^ **(Fig. 4b**), suggesting that AZ localization may be precisely predicted from the localization of the PSD scaffold. In order to verify this empirically, we measured the length, width, and depth of Homer1 and Bassoon (a major AZ scaffolding protein) appositions (Supp. Methods). All parameters were significantly correlated between corresponding appositions (n=140 over 3 independent experiments; Spearman’s R for length: r=0.61, p<0.0001; width: r=0.55, p<0.0001; depth: r=0.34, p<0.0003) and the approximately 0.1 μm^2^ mean area size of individual appositions (see Suppl. Methods) corresponds to the recently reported size of individual nanomodules^17^. Furthermore, distances between Homer1 and Bassoon protein clusters varied very little between synapses and were not affected under CB_1_R activation (**Fig. 4c-e**). Based on these properties, we were successfully able to predict the 3D location of the presynaptic AZ volume (AZv, **Fig. 4f**) based on postsynaptic Homer1 clusters. Indeed, on an independent sample 95% of Bassoon locations were contained within our predicted AZv (**Fig. 4g**).

**Figure 4:**
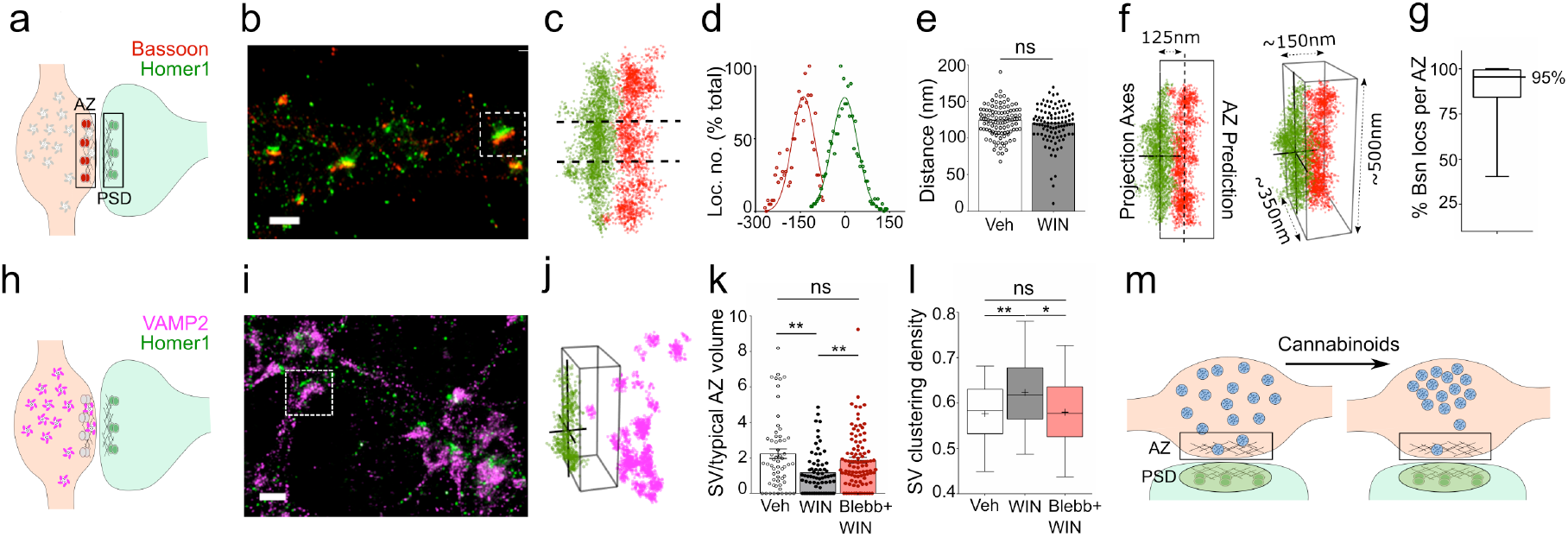
3D STORM reveals synaptic vesicle redistribution with actomyosin contraction under CB_1_R activation. **(a)** Representation of Bassoon and Homer1 distributions within the pre-synaptic active zone (AZ) and post-synaptic density (PSD), respectively. (**b)** STORM reconstruction of Bassoon and Homer1 localizations **(c)**. Boxed synapse in (b) after clustering analysis. Dotted lines enclose the localisations averaged over the x-axis for Gaussian fitting in *d*. (**d)** Averaged localisations and Gaussian fit of synapse in *c*. Distance between Gaussian peaks represents the measured distance between Homer1 and Bassoon appositions. (**e)** Measured distances between pre- and post-synaptic appositions under control conditions (Veh) and CB_1_R activation (WIN). (Veh: n=105, 122.3±2.0nm, WIN: n=100, 117.9±2.6nm; p=0.361, Mann-Whitney test). (**f)** Prediction of the AZ based on both PSD axes and the median apposition distances measured in *e*. Side values in the 3D view (right) represent average measures obtained from predictions over 105 synapses for the AZ volume (AZv). (**g**) Percentage of Bassoon (Bsn) localisations per synapse contained within the predicted AZv in an independent sample (n=55, 25^th^percentile: 84.3%; 75^th^percentile: 99.4%). (**h**) Representation of Vamp2 and Homer1 distributions within the synapse. (**i**). STORM reconstruction of Vamp2 and Homer1 (**j**) Synapse boxed in (h) after identification of synaptic vesicles through nested clustering analysis. The 3D box shows the predicted AZv for this synapse, enclosing one identified vesicle. (**k**) Fewer vesicles are found within the predicted AZv under CB_1_R activation (Veh: 87.8±10.6/μm^3^, n=59 over 3 independent experiments; WIN: 45.4±5.5/μm^3^, n=70 over 3 independent experiments, p<0.003). This depletion is prevented by inhibiting actomyosin contraction with para-nitroblebbistatin (Blebb) (73.2±6.2/μm^3^, n=98 over 3 independent experiments, p<0.004). Values are normalized to a typical AZv (=2.5×10^7^nm^3^). (**i**) Vesicle clustering found under CB_1_R activation is prevented by blocking actomyosin contraction. (**m**) Putative mechanisms of CB_1_R activation on vesicle redistribution during CB_1_R-induced plasticity. Scale bars in *b* and *i*: 1μm. Kruskal-Wallis test in *k* and *l*.

We next immunolabeled synaptic vesicles for the SNARE protein VAMP2 (**Fig. 4h**), and developed a nested clustering algorithm to identify synaptic vesicles within VAMP2 localisation clusters (**Fig.4j**; Supp. Methods), to assess the spatial organization of the synaptic vesicles pool. We found strikingly fewer synaptic vesicles in the predicted AZv under WIN55,212-2 treatment (1μM, 10min), whereas pretreatment with para-nitroblebbistatin (25μM, 20min) significantly prevented this effect. **(Fig.4k)**. Furthermore, synaptic vesicles were significantly more clustered to each other within the total pool under WIN55,212-2 treatment compared to either control conditions or para-nitroblebbistatin pretreatment **(Fig.4l)**. These results imply that CB_1_R activation induces a significant redistribution of synaptic vesicles within the presynaptic compartment following CB_1_R activation, leading to depletion of synaptic vesicles from the AZ. Inhibition of actomyosin contraction prevents both CB_1_R-induced vesicle redistribution and presynaptic silencing, suggesting that cannabinoid-induced changes in synaptic efficiency depend on the contractile properties of the presynaptic actomyosin cytoskeleton **(Fig.4m).**

In conclusion, in this study we have combined patch-clamp recordings with super resolution microscopy and functional imaging to establish the link between presynaptic architecture and synapse function at two archetypal CNS synapses. By showing that CB_1_R dynamically controls presynaptic organization associated to individual synaptic nanomodules^17^ through actomyosin contractility, our results provide both a mechanism and a functional relevance to recent electron microscopy studies that reported fewer synaptic vesicles near the presynaptic active zone following cannabinoid treatment both *in vitro* and *in vivo* ^13,19^. Our results significantly extend the suggested roles for the presynaptic actin cytoskeleton, hopefully advancing toward a more complete model of regulation of vesicle release^20^. Structurally, synaptic vesicles are reversibly tethered to the actin cytoskeleton by synapsin, particularly vesicles of the recycling and/or resting pool ^21^. Actin cytoskeleton dynamics also have an important role in the preferential distribution of recycling vesicles close to the AZ^22^. Our findings now identify another major actin binding/structuring protein, the contractile NMII, mechanistically explaining through dynamic redistribution of vesicles, the generation of a widespread form of long-term presynaptic plasticity. Notably, we did not find an effect of either NMII or ROCK inhibition on short-term forms of eCB-induced synaptic plasticity, neither on DSI nor DSE. Indeed, eCB-STD occur very rapidly (<1sec), following brief activation of CB_1_Rs, possibly effecting only already docked vesicles, while a longer CB_1_R activation (>5min) is necessary for eCB-LTD induction^23^. This longer activation period may be needed to engage actomyosin contraction, a molecular mechanism leading to actin cytoskeleton remodeling over several minutes in non-muscular cells^24^. Our results, by reporting a conceptually novel molecular mechanism of synaptic plasticity downstream of an important recreational drug target and known risk factor in schizophrenia ^25^ open novel perspectives in the understanding of cognitive function, the pathogenesis of marijuana intoxication and neuro-psychiatric disease.

## Methods

### Animals

Experiments were performed in accordance with local animal welfare committee (Center for Interdisciplinary Research in Biology and EU guidelines; directive 2010/63/EU). Rats and mice (Charles River, L’Arbresle, France) were housed in standard 12 hours light/dark cycles and food and water were available *ad libitum*.

### Antibodies and Chemicals

Rabbit polyclonal Homer1 (Cat. No. 160 003) and mouse monoclonal VAMP2 (Cat. No. 104 211) antibodies were obtained from Synaptic Systems (Goettingen, Germany). Bassoon mouse monoclonal antibody (Cat. No. ab82958) was obtained from Abcam (Paris, France). Paired fluorophore-conjugated secondary antibodies were made as previously described^26^. WIN55,212-2 and (RS)-3,5-DHPG were from Tocris. Carbachol, Y-27632, active (S)-(−)-blebbistatin and inactive (R)-(+)-blebbistatin enantiomers and para-nitroblebbistatin were from Calbiochem, Sigma and Optopharma. None of the bath-applied drugs had a significant effect on basal IPSC and EPSC amplitudes, in our experimental conditions.

### Time-lapse microscopy of primary cultured neurons

Dissociated neurons obtained from hippocampi of day 17-18 Sprague-Dawley rat embryos were plated on Poly-D-Lysine-coated coverslips at a density of approximately 100,000 cells per coverslip and subsequently cultivated at 37°C, 5% CO_2_ in Neurobasal^TM^ (LifeTech) medium supplemented with 2% B27 (LifeTech), 0.5mM L-glutamine, 10U/mL penicillin G and 10mg/mL streptomycin containing conditioned medium, obtained by incubation with glial cultures (70-80% confluence) for 24 h as described previously^5^. Neurons were transfected either with Synaptophysin-pHluorin (SpH), a kind gift from Dr. Stefan Krueger (Dalhousie University, Halifax, NS, Canada), and LifeAct-mCherry^5^, or with SpH alone. Transfections and time-lapse microscopy were performed at 37°C, as described previously^5^. Transfections were performed after 7-9 days *in vitro* (DIV), and time-lapse microscopy was performed after 17-21 DIV. Briefly, coverslips were placed in a Ludin chamber (Life Imaging Services, Basel, Switzerland) filled with imaging buffer (120 mM NaCl, 3 mM KCl, 2 mM CaCl2, 2 mM MgCl2, 10 mM glucose, and 10 mM HEPES, pH 7.35). Neurons were depolarized by adding 50mM KCl for 2min, followed by NH_4_Cl treatment. Pretreatments and treatments were applied at 30min and 10min before KCl, respectively. Dimethylsulfoxide vehicle concentrations ranged from 0.02% to 0.1%.

Image stacks were realigned using ImageJ. SpH fluorescence intensity was measured in round ROIs of approximately 3×3μm, placed manually around visually identified axonal boutons, with mean basal axonal fluorescence intensity subtracted for each timepoint. Axonal boutons were selected for analysis if SpH response to NH_4_Cl was superior to that of KCL and if baseline fluorescence was within 2x the standard deviation around baseline population mean. Statistical analyses used Kruskal-Wallis test with Dunn’s post-hoc Multiple Comparison Test, n indicates the number of axonal boutons analyzed. Values are mean± SEM.

### Electrophysiological recordings and analysis from hippocampal slices

400 μM transverse hippocampal vibratome slices were prepared from 6- to 8-week-old C57BL6 male mice in ice-cold extracellular solution containing (in mM): 10 NaCl, 195 sucrose, 2.5 KCl, 15 glucose, 26 NaHCO_3_, 1.25 NaH_2_PO_4_, 1 CaCl_2_ and 2 MgCl_2_). The slices were then transferred to 30°C ACSF (in mM: 125 NaCl, 2.5 KCl, 10 glucose, 26 NaHCO_3_, 1.25 NaH_2_PO_4_, 2 Na Pyruvate, 2 CaCl_2_ and 1 MgCl_2_) for 30min and kept at room temperature for at least 1.5 hours before recording at 33°C. Cutting and recording solutions were both saturated with 95% O2 and 5% CO2 (pH 7.4).

Whole-cell recordings were obtained using Axograph X software from CA1 PNs in voltage clamp mode in the continuous presence of the NMDA receptor antagonist d-(-)-2-amino-5-phosphonopentanoic acid (d-APV; 50 μM) and the AMPA/kainate receptor antagonist 2,3-dihydroxy-6-nitro-7-sulfonyl-benzo[f]quinoxaline (NBQX; 10 μm) Inhibitory currents were recorded at +10 mV with a patch pipette (3–5 MΩ) containing (in mM): 135 CsMethylSulfate, 5 KCl, 0.1 EGTA-Na, 10 HEPES, 2 NaCl, 5 ATP, 0.4 GTP, 10 phosphocreatine (pH 7.2; 280–290 mOsm). Series resistance (typically 12–18MΩ) was monitored throughout each experiment; cells with more than 15% change in series resistance were excluded from analysis. Synaptic potentials were evoked by monopolar stimulation with a patch pipette filled with ACSF and positioned in the middle of CA1 SR. A HFS (100 pulses at 100Hz repeated twice) was applied following 15 – 20min of stable baseline. The amplitudes of the IPSCs were normalized to the baseline amplitude. The magnitude of LTD was estimated by comparing averaged responses at 30-40min after the induction protocol with baseline-averaged responses 0–10min before the induction protocol.

### Electrophysiological recordings and analysis from corticostriatal slices

330μm horizontal brain slices containing the somatosensory cortex and the corresponding corticostriatal projection field in the dorsolateral striatum were prepared from P_25-35_ male rats as previously described^12,27^. Brains were sliced in a 95% CO2/5% O2-bubbled, ice-cold cutting solution containing (in mM) 125 NaCl, 2.5 KCl, 25 glucose, 25 NaHCO_3_, 1.25 NaH_2_PO_4_, 2 CaCl_2_, 1 MgCl_2_, 1 pyruvic acid, and then transferred into the same solution at 34°C for one hour and then moved to room temperature before patch-clamp whole-cell recordings (at 34°C).

Patch-clamp recordings were performed as previously described^12,27^. Briefly, borosilicate glass pipettes of 4-6MOhms resistance contained for whole-cell recordings (in mM): 105 K-gluconate, 30 KCl, 10 HEPES, 10 phosphocreatine, 4 ATP-Mg, 0.3 GTP-Na, 0.3 EGTA (adjusted to pH 7.35 with KOH). The composition of the extracellular solution was (mM): 125 NaCl, 2.5 KCl, 25 glucose, 25 NaHCO_3_, 1.25 NaH_2_PO_4_, 2 CaCl_2_, 1 MgCl_2_, 10mM pyruvic acid bubbled with 95% O_2_ and 5% CO_2_. Signals were amplified using EPC10-2 amplifier (HEKA Elektronik, Lambrecht, Germany). All recordings were performed at 34°C and slices were continuously superfused at 2-3ml/min with the extracellular solution. Slices were visualized on an Olympus BX51WI microscope (Olympus, Rungis, France) using a 4x/0.13 objective for the placement of the stimulating electrode and a 40x/0.80 water-immersion objective for localizing cells for whole-cell recordings. Series resistance was not compensated. Current-clamp recordings were filtered at 2.5kHz and sampled at 5kHz and voltage-clamp recordings were filtered at 5kHz and sampled at 10kHz using the Patchmaster v2×32 program (HEKA Elektronik). Electrical stimulations were performed with a bipolar electrode (Phymep) placed in the layer 5 of the somatosensory cortex and were monophasic at constant current (ISO-Flex stimulator)^10,12,27^. Currents were adjusted to evoke striatal EPSCs ranging in amplitude from 50 to 200pA. Repetitive control stimuli were applied at a frequency of 0.1Hz for 60min after LFS protocol. Recordings on neurons were made over a period of 10 minutes at baseline, and for at least 60 minutes after the LFS protocols; long-term changes in synaptic efficacy were measured from 45 to 55 minutes. We individually measured and averaged 60 successive EPSCs, comparing the last 10 minutes of the recording with the 10-minute baseline recording. Series resistance was monitored for each sweep and a variation above 20% led to the rejection of the experiment. LTD was induced with low frequency stimulation protocol consisting in 600 cortical stimulations at 1Hz paired with postsynaptic concomitant depolarization of the MSN during 50ms^10,12^. For DSE induction, MSN was depolarized from RMP to 0mV during 10sec (with bath-applied carbachol, 10μM, and DHPG, 50μM)^10^. Off-line analysis was performed using Fitmaster (Heka Elektronik) and Igor-Pro 6.0.3 (Wavemetrics, Lake Oswego, OR, USA). Statistical analysis was performed using Prism 5.0 software (San Diego, CA, USA). “n” refers to a single cell experiment from a single slice.

### Immunocytochemistry and STORM imaging

Neurons used for STORM imaging underwent the same treatment protocol as for videomicroscopy at 21-28 DIV, with the exception that instead of depolarization with KCl after treatment, neurons were fixed with a preheated solution of 4% PFA and 4% sucrose in 0.1 M phosphate buffered saline (PBS) for 15 minutes at room temperature (RT), permeabilized after wash for 5 min at RT with 0.1% Triton X in PBS, and blocked for 1h at RT with blocking buffer (4% BSA in PBS). Primary and secondary antibodies were applied in blocking buffer, primary antibodies being applied overnight at 4°C and secondary antibodies being applied for 2h at RT. After washing out the secondary antibody, neurons were post-fixed with 4% PFA and 4% sucrose in PBS for 5min, washed and stored in PBS at 4°C before imaging.

STORM images were acquired on a N-STORM microscope (Nikon Instruments), outfitted with 405 nm, 561 nm, and 647 nm solid-state lasers, a 100X NA 1.49 objective and an Ixon DU-897 camera. Imaging was performed as previously described^26^. Briefly, visually identified dendrites labelled with activator-reporter fluorophore pairs (Alexa Fluor 405 –Alexa Fluor 647 and Cy3-Alexa Fluor 647) were imaged using sequences of one activator frame (405 or 561 nm) followed by three reporter frames (647 nm)^16^. A cylindrical lens was placed across the optical path in order to acquire 3D information^27^, and the N-STORM software (Nikon Instruments) was used for the localization of single fluorophores.

## Acknowledgements

This work was supported by the MyoSynapse grant of Paris Sciences et Lettres Research University and the Ecole des Neurosciences de Paris.

**Author Contributions** *MHM, CL, LV, VC and LZ designed research, MHM, HX, YC, SP, RAP, AMC, DZ and VC realized the experiments, MHM, RAP, LV, VC and LZ wrote the article.*

## References

1. Ohno-Shosaku, T. & Kano, M. Endocannabinoid-mediated retrograde modulation of synaptic transmission. Current Opinion in Neurobiology 29C, 1–8 (2014).

2. Wilson, R. I. & Nicoll, R. A. Endogenous cannabinoids mediate retrograde signalling at hippocampal synapses. Nature 410, 588–592 (2001).

3. Castillo, P. E., Younts, T. J., Chávez, A. E. & Hashimotodani, Y. Endocannabinoid Signaling and Synaptic Function. Neuron 76, 70–81 (2012).

4. Monday, H. R. & Castillo, P. E. Closing the gap: long-term presynaptic plasticity in brain function and disease. Current Opinion in Neurobiology 45, 106–112 (2017).

5. Roland, A. B. et al. Cannabinoid-induced actomyosin contractility shapes neuronal morphology and growth. Elife 3, e03159–e03159 (2014).

6. Holtmaat, A., Randall, J. & Cane, M. Optical imaging of structural and functional synaptic plasticity in vivo. European Journal of Pharmacology 719, 128–136 (2013).

7. Kovacs, M., Toth, J., Hetenyi, C., Malnasi-Csizmadia, A. & Sellers, J. R. Mechanism of blebbistatin inhibition of myosin II. Journal of Biological Chemistry 279, 35557–35563 (2004).

8. Heifets, B. D. & Castillo, P. E. Endocannabinoid Signaling and Long-Term Synaptic Plasticity. Annu. Rev. Physiol. 71, 283–306 (2009).

9. Chevaleyre, V. et al. Endocannabinoid-mediated long-term plasticity requires cAMP/PKA signaling and RIM1alpha. Neuron 54, 801–812 (2007).

10. Puente, N. et al. Polymodal activation of the endocannabinoid system in the extended amygdala. Nature Neuroscience 14, 1542–1547 (2011).

11. Atwood, B. K. & Lovinger, D. M. in Endocannabinoids and Lipid Mediators in Brain Functions 27, 109–153 (Springer International Publishing, 2017).

12. Fino, E., Glowinski, J. & Venance, L. Bidirectional activity-dependent plasticity at corticostriatal synapses. Journal of Neuroscience 25, 11279–11287 (2005).

13. Ramírez-Franco, J., Bartolomé-Martín, D., Alonso, B., Torres, M. & Sánchez-Prieto, J. Cannabinoid type 1 receptors transiently silence glutamatergic nerve terminals of cultured cerebellar granule cells. PLoS ONE 9, e88594–e88594 (2013).

14. Miesenböck, G., De Angelis, D. A. & Rothman, J. E. Visualizing secretion and synaptic transmission with pH-sensitive green fluorescent proteins. Nature 394, 192–195 (1998).

15. Képiró, M. et al. para-Nitroblebbistatin, the Non-Cytotoxic and Photostable Myosin II Inhibitor. Angew. Chem. Int. Ed. 53, 8211–8215 (2014).

16. Bates, M., Huang, B., Dempsey, G. T. & Zhuang, X. Multicolor super-resolution imaging with photo-switchable fluorescent probes. Science 317, 1749–1753 (2007).

17. Hruska, M., Henderson, N., Le Marchand, S. J., Jafri, H. & Dalva, M. B. Synaptic nanomodules underlie the organization and plasticity of spine synapses. Nature Neuroscience 21, 671–682 (2018).

18. Tang, A.-H. et al. A trans-synaptic nanocolumn aligns neurotransmitter release to receptors. Nature 536, 210–214 (2016).

19. García-Morales, V., Montero, F. & Moreno-López, B. Cannabinoid agonists rearrange synaptic vesicles at excitatory synapses and depress motoneuron activity in vivo. Neuropharmacology (2015).

20. Rust, M. B. & Maritzen, T. Relevance of presynaptic actin dynamics for synapse function and mouse behavior. Experimental Cell Research 335, 165–171 (2015).

21. Cesca, F., Baldelli, P., Valtorta, F. & Benfenati, F. The synapsins: key actors of synapse function and plasticity. Progress in Neurobiology 91, 313–348 (2010).

22. Marra, V. et al. A Preferentially Segregated Recycling Vesicle Pool of Limited Size Supports Neurotransmission in Native Central Synapses. Neuron 76, 579–589 (2012).

23. Chevaleyre, V., Takahashi, K. A. & Castillo, P. E. Endocannabinoid-mediated synaptic plasticity in the CNS. Annu. Rev. Neurosci. 29, 37–76 (2006).

24. Li, D. et al. ADVANCED IMAGING. Extended-resolution structured illumination imaging of endocytic and cytoskeletal dynamics. Science 349, aab3500–aab3500 (2015).

25. Large, M. The need for health warnings about cannabis and psychosis. - PubMed - NCBI. The Lancet Psychiatry 3, 188–189 (2016).

26. Leterrier, C. et al. Nanoscale Architecture of the Axon Initial Segment Reveals an Organized and Robust Scaffold. Cell Rep 13, 2781–2793 (2015).

27. Cui, Y. et al. Endocannabinoid dynamics gate spike-timing dependent depression and potentiation. Elife 5, e13185 (2016).

27. Huang, B., Wang, W., Bates, M. & Zhuang, X. Three-dimensional super-resolution imaging by stochastic optical reconstruction microscopy. Science 319, 810–3 (2008).

